# *Caenorhabditis elegans* MES-3 is a highly divergent ortholog of the canonical PRC2 component SUZ12

**DOI:** 10.1101/2022.04.07.487461

**Authors:** Berend Snel, Sander van den Heuvel, Michael F. Seidl

## Abstract

Polycomb Repressive Complex 2 (PRC2) catalyzes the mono-, di, and trimethylation of histone protein H3 on lysine 27 (H3K27), which is strongly associated with transcriptionally silent chromatin. The functional core of PRC2 is highly conserved in animals and consists of four subunits. One of these, SUZ12, has not been identified in the genetic model *Caenorhabditis elegans*, whereas *C. elegans* PRC2 contains the clade-specific MES-3 protein. Through unbiased sensitive sequence similarity searches complemented by high-quality structure predictions of monomers and multimers, we here demonstrate that MES-3 is a highly divergent ortholog of SUZ12. MES-3 shares protein folds and conserved residues of key domains with SUZ12 and is predicted to interact with core PRC2 members similar to SUZ12 in human PRC2. Thus, in agreement with previous genetic and biochemical studies, we provide evidence that *C. elegans* contains a diverged yet evolutionary conserved core PRC2, like other animals.

## INTRODUCTION

Post-translational modifications of histone proteins contribute to the organization of genomic DNA and establishment of transcriptionally active versus silent chromatin^1^. Polycomb group proteins form an important class of transcriptional repressors that function through modification of histone tails^2,3^. These proteins assemble into two distinct multi-subunit complexes, Polycomb Repressive Complex 1 and 2 (PRC1 and PRC2)^1–5^. PRC2 catalyzes the mono-, di-, and trimethylation of histone protein H3 on lysine 27 (H3K27), which is strongly associated with transcriptionally silent chromatin and plays an important role in the maintenance of cell identity and developmental regulation gene expression.

The functional core of PRC2 is highly conserved in animals and consists of four subunits: the H2K27 methyltransferase EZH2/1 and associated proteins EED, SUZ12, and RBBP4/7^4–6^ (**Fig. 1a, b)**. SUZ12 interacts with all members of the PRC2 core to form two distinct lobes^6–8^. The N-terminal region of SUZ12 together with RBBP4/7 forms the targeting lobe, which contributes to the recruitment and regulation of PRC2, and serves as a platform for co-factor binding^7,8^. The region of SUZ12 included in this lobe contains five motifs and domains: the zinc finger binding (ZnB), WD-domain binding 1 (WDB1), C2 domain, zinc finger (Zn), and WD-domain binding 2 (WDB2)^7,8^ (**Fig. 1b**). The C-terminal region of SUZ12 contains a VEFS domain (**Fig. 1b**), which associates with EZH2/1 and EED to form the catalytic lobe of PRC2^7,8^. Thus, SUZ12 is critical for the assembly, integrity, and function of PRC2, in agreement with the conservation of SUZ12 as a core PRC2 component in animals (**Fig. 1a**).

**Figure 1.**
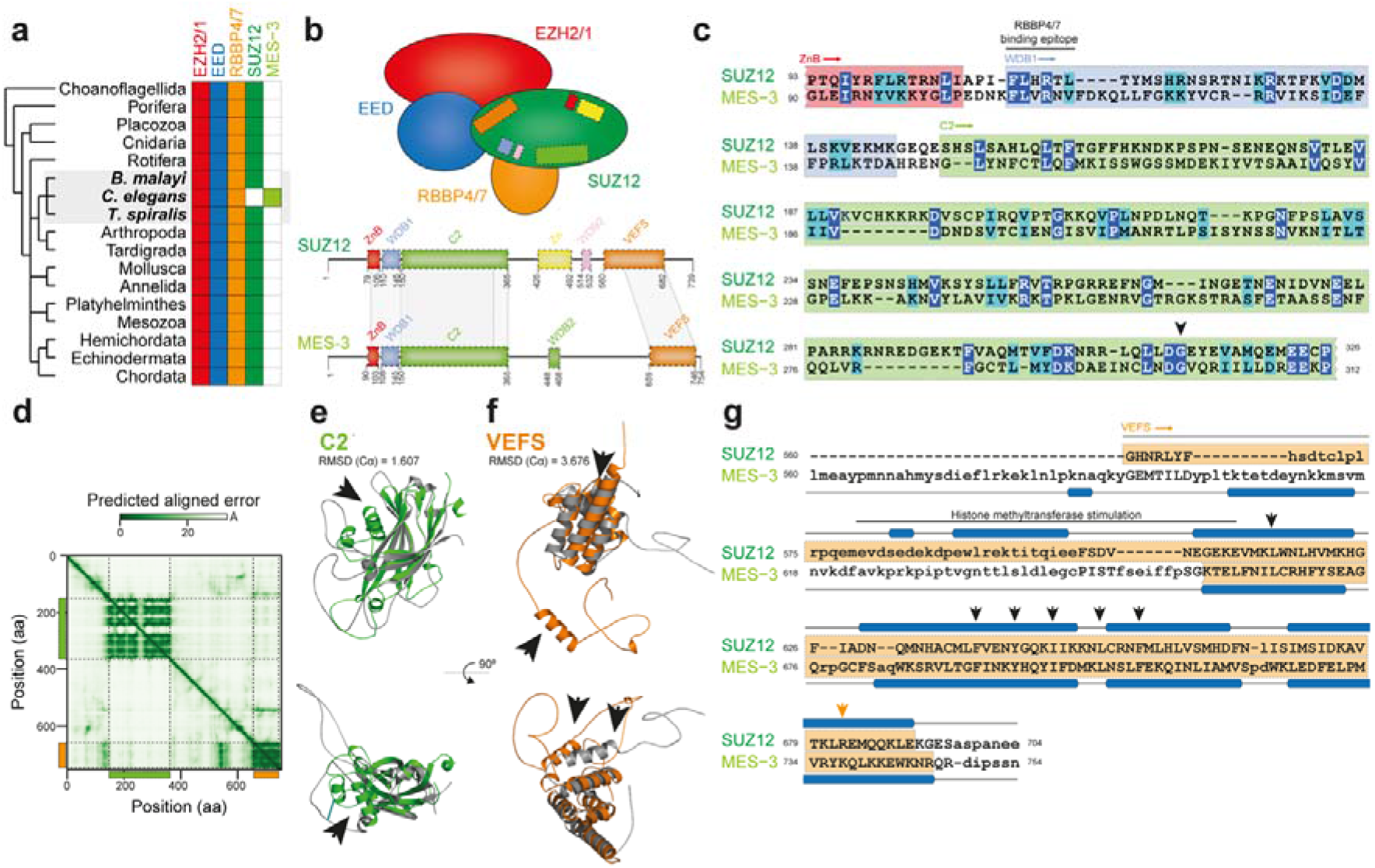
MES-3 is a highly divergent ortholog of the canonical Polycomb Repressive Complex 2 component SUZ12. **a**. The Polycomb Repressive Complex 2 (PRC2) core components EZH2/1, EED, RBBP4/7, and SUZ12 are conserved in a broad range of metazoans; the presence of orthologs is indicated by filled boxes. Notably, based on sequence similarity searches, an ortholog of SUZ12 is absent in the nematode model species *Caenorhabditis elegans*, but present in other, closely related nematodes (*Brugia malayi* and *Trichinella spiralis*). *C. elegans* encodes the PRC2 core component MES-3^9,17^ that lack obvious motifs or sequence similarity to SUZ12^17^. **b**. Schematic representation of the composition of the core PRC2. The zinc finger binding (ZnB; red), WD-domain binding 1 (WDB1; blue), C2 domain (green), zinc finger (Zn; yellow), WD-domain binding 2 (WDB2; pink), and VEFS (orange) motifs or domains involved in SUZ12 protein-interactions are shown in the schematic as well as along the protein sequence^7,8,26^. Schematic representation of the protein sequence of MES-3 is shown, and regions of here uncovered sequence (**c**) and structural (**e, f**) similarity are highlighted. **c**. Protein sequence alignment between the N-terminal region of SUZ12 and MES-3, as identified by sensitive profile-vs-profile sequence similarity searches, covers part of the zinc finger binding (ZnB; red), WD-domain binding 1 (WDB1; blue), and C2 domain (green). The conserved RBBP4/7 binding epitope as well as Gly299 are highlighted^19–21^. Identical amino acids are shown in blue and biochemically similar amino acids are shown in turquoise. **d**. The predicted aligned error (in Å; based on model 2 ptm) of the MES-3 structure is shown as a heatmap and reveals two separated globular regions in the N- and C-terminus, the former overlaps with the profile-vs-profile match (**c**) and corresponds to the C2 domain of SUZ12 (**e; Fig. S1i;** RMSD = 1.607), while the latter overlaps with the regions that structurally resemble the VEFS domain (**f; Fig. S1j;** RMSD = 3.676). The black arrows (**e, f**) highlight regions that differ considerably between SUZ12 and MES-3 (**Fig. S1i, j**), and the structure predictions of SUZ12 and MES-3 (**e, f**) are shown in grey as well as green (C2) and orange (VEFS), respectively. **g**. Sequence-independent structure alignment of the VEFS regions of SUZ12 and MES-3 reveals significantly structural similarity (Dali Z-score = 8.3; TM-score = 0.55), especially along the alpha helices in the C-terminus; a region previously shown to stimulate histone methyltransferase activity in SUZ12^20^ (pos. 580 to 612) is highlighted by a black bar, and individual amino acids important for PRC2 assembly^20^ are shown by black arrows.

Genetic and biochemical studies in the model nematode *Caenorhabditis elegans* revealed a functional PRC2 complex without an apparent SUZ12 ortholog^9–15^. The components of this complex were originally defined by specific maternal-effect sterile (*mes*) mutations that cause defects in germline development and silencing of the X chromosome in the hermaphrodite germline^10,16^. Molecular characterizations revealed that MES-2 and MES-6 are homologs of the Polycomb group proteins EZH2/1 and EED, respectively^9^. MES-2 (EZH2/1) and MES-6 (EED) form a protein complex with MES-3, and all three components are required for histone H3 K27 methyltransferase activity *in vivo* and *in vitro*^9,11–14^. Despite the functional similarity with the PRC2 core, MES-3 appeared to lack obvious motifs or sequence similarity to SUZ12 or RBBP4/7, and therefore has been considered a *C. elegans* specific subunit^9,12,13,15,17^. Consequently, PRC2 in *C. elegans* and in animals are considered functional analogues, despite a seemingly divergent subunit composition^9,12,13,15^. In-depth sequence comparisons have recently turned up surprising homologies^18^, which prompted us to investigate whether MES-3 could be a highly diverged homolog of SUZ12 instead of a *C. elegans* specific invention.

## RESULTS & DISCUSSION

To identify MES-3 homologs in animals, we used unbiased sensitive profile-vs-profile searches to query the predicted human proteome with MES-3 and query the worm proteome with SUZ12. Surprisingly, we recovered a consistent but insignificant bidirectional match between SUZ12 and MES-3 (16% identity; **Fig. 1c**) that is located at approximately the same regions in both proteins and covers 223 amino acids in MES-3. This region in SUZ12 spans part of the ZnB motif, the complete WDB1 motif, and most of the C2 domain (**Fig. 1b, c**). Notably, the conserved RBBP4/7 binding site of SUZ12^19^ is also present in MES-3 (pos. 108-113; FLxRx[VL]) as well as a conserved glycine (pos. 299) (**Fig. 1c**); a missense mutation of this glycine in *Drosophila* leads to a partial loss-of-function phenotype^20,21^. Therefore, we conclude that the N-terminal region of SUZ12 and MES-3 shares extended sequence similarity including residues previously shown to be critical for function, suggesting that these two proteins are homologs. However, the profile-to-profile searches did not detect similarity between the C-terminal sequence of MES-3 and the SUZ12 domain that mediates EZH2 and EED interaction^7,8^ (**Fig. 1b**).

Protein structure is typically more conserved than primary sequence and better allows detection of diverged homologs^22^. Since the protein structure of MES-3 is not yet experimentally resolved, we used deep-learning driven protein structure prediction of both MES-3 and SUZ12 with Alphafold2^23^. The SUZ12 structure has six functional motifs and domains that were predicted with high precision as they resemble the experimentally determined structure (RMSD = 0.56-1.14; global TM-score = 0.70; global Dali Z-score = 14.8 **Fig. S1a-e**). Like SUZ12, the predicted MES-3 structure is partially disordered (**Fig. 1d; S1f-h**), but nevertheless has a globular N-terminal region mainly formed by β-sheets and a C-terminal region mainly formed by α-helices (**Fig. 1d, e**), and both regions were modelled with high confidence (**Fig. S1g**). Interestingly, the C2 domain of SUZ12 shares significant structural similarity with the N-terminal structural regions of MES-3 (**Fig. 1d,e; Fig. S1i**; RMSD = 1.607; TM-score = 0.60; Dali Z-score = 11.6), corroborating the profile-vs-profile results (**Fig. 1c**). The structural similarity (MES-3, pos. 150-365) extends beyond the region of shared sequence similarity identified above (MES-3, pos. 150-312), and thus encompasses the complete C2 domain (**Fig. 1d; Fig. S1i**). Nevertheless, we also observed some differences in the predicted structures such as the occurrence of an unmatched alpha helix in MES-3 (**Fig. 1e; Fig. S1i**) or the absence of amino acids in MES-3 known to be involved in the interaction between SUZ12 and RBBP4/7 (e.g., SUZ12, R196^8^).

Likewise, we observed structural similarity between the C-terminal domain of MES-3 and the VEFS domain in SUZ12 (**Fig. 1b, d, f; g;** RMSD = 3.676; TM-score = 0.55; Dali Z-score = 8.3). The MES-3 VEFS-like region is considerably shorter compared with SUZ12 and lacks amino acids that are thought to be involved in the stimulation of histone methyltransferase activity (SUZ12, pos. 580 to 612^21^) and specifically SUZ12 E610 and K611^21^, which are invariant in plants, animals, and fungi (**Fig. 1g**; **Fig. S1j**). By contrast, several bulky or hydrophobic aromatic residues whose deletion impacts PRC2 assembly^20,21^ are conserved, e.g., SUZ12 pos. F639, I647, L652, and F656 can be aligned to identical residues in superposition of the SUZ12 and MES3-VEFS predicted structures (**Fig. 1g**; **Fig. S1j**). This suggests that even though the overall sequence similarity is very low, the VEFS domain is overall well conserved in MES-3.

MES-3 together with MES-2 (EZH2) and MES-6 (EED) forms a stable heterotrimeric protein complex^9,15^. To identify potential interaction surfaces of MES-3, we used Alphafold2^23,24^ to generate high-quality structure predictions for MES-2 and MES-6 monomers (**Fig. S2a-l**) as well as the trimeric MES-2, MES-3, and MES-6 core complex (**Fig. 2a, c; Fig S2m**). As in human PRC2^7,8,25,26^ (**Fig. 2b**), the C-terminal VEFS domain of MES-3 is predicted to be associated with MES-2 and MES-3 (**Fig. 2a, c, f**). Even though the VEFS domain in MES-3 is shorter than in SUZ12^26^ (**Fig. 1g**), it interacts with a region of MES-2 (pos. 300 to 450; **Fig. 2c, f**) that in EZH2 comprises the MCSS and the SANT2 domain, which together with VEFS stimulate histone methyltransferase activity^21,26^. While these elements were previously noted to be absent in MES-2^15^, our comparison suggests that this region in MES-2 shows potentially similar structural elements as well as considerable sequence divergence compared with EZH2. We also identified a short region of MES-3 (pos. 530-570) that is associated with regions in both MES-2 and MES-6 (**Fig. 2c, f**). The N-terminal region of SUZ12 together with RBBP4/7 forms the targeting lobe^8,25,26^, and thus we sought to predict interaction surfaces between MES-3 and LIN-53, one of two closely related seven WD40-repeat proteins, and the protein that most likely retained the ancestral RBBP4/7 function (**Fig. 2d**; **Fig S2n**). Similar to human PRC2^8,25,26^ (**Fig. 2b**), we observed interactions of the WDB1 domain with the WD40 repeats at the N- and C-terminus of LIN-53 (**Fig. 2e, f**). We also identified a second short region in MES-3 (pos. 448-468) that interacts with N-terminal WD40 repeats in LIN-53, resembling the interaction of WDB2 in human PRC2^8,25^, and thus MES-3 WDB1 and WDB2 wrap around WD40 repeats of LIN-53 (**Fig. 2f**), which in human PRC2 inhibits H3K4 binding of RBBP4/7^8^.

**Figure 2.**
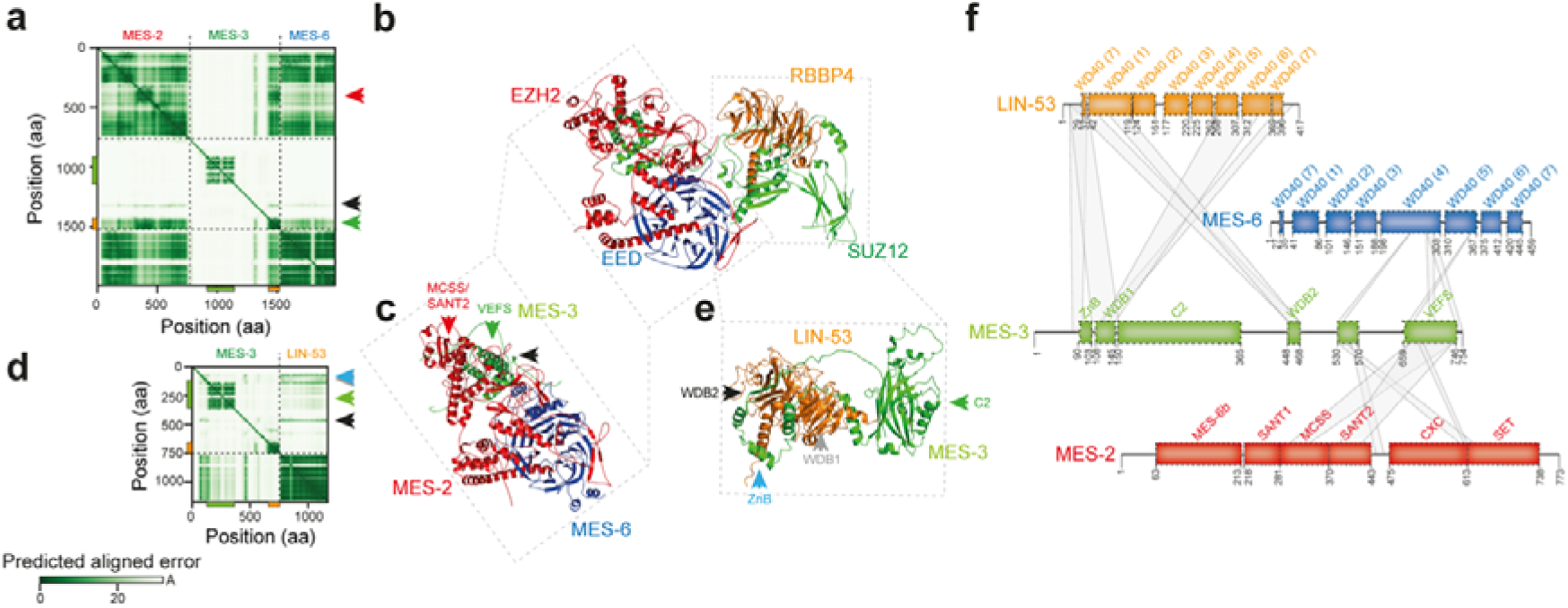
MES-3 provides a structural framework for PRC2 in *C. elegans*. **a**. The predicted aligned error (in Å) of MES-3 co-folded with MES-2 and MES-6 is shown as a heatmap and is consistent with association of MES-3 with MES-2 and MES-6 in the C-terminal regions of MES-3, which overlaps with the predicted VEFS domain. **b**. Experimentally resolved human core PRC2 (6WKR^25^) highlights interactions between SUZ12 and RBBP4 as well as SUZ12 and EZH2 and EED. **c**. Predicted *C. elegans* core PRC2 is formed by MES2, MES-3, and MES-6. The corresponding region in human PRC2 is highlighted, as well as the position of the MES-3 VEFS domain (green triangle, see **a**.) and the MES-2 MCSS/SANT2 region (red triangle, see **a**.), as well as a short central region of MES-3 (pos. 530-570) that associates with multiple regions in MES-2 and MES-6 (black triangle, see **a**.). For clarity, only regions of MES-3 interacting with MES-2 and MES-6 are shown (pos. 1-530 and 570-640 are hidden). **d**. The predicted aligned error (in Å) of MES-3 co-folded with LIN-53 is shown as a heatmap and reveals association between the N-terminal region of MES-3 and LIN-53. **e**. Predicted MES-3 with LIN-53 complex. The corresponding region in human PRC2 is highlighted, as well as the MES-3 C2 domain (green triangle, see **d**.) and regions surrounding the C2 domain that engage in association with LIN-53 (WDB2, black triangle; WDB1, grey triangle; ZnB, light-blue triangle; see **d**.). For clarity, only regions of MES-3 interacting with LIN-53 are shown (pos. 1-80, 365-415, and 470-754 are hidden). **f**. Schematic representation of MES-3 and its predicted interactions with MES-2, MES-6, and LIN-53. The positions of protein domains/motifs in MES-2, MES-6, and LIN-53 were inferred via structural alignments of the predicted structures to those of the experimentally determined human EZH2, EED, and RBBP4^25,26^. MES-3 domains/motifs are indicated as in **Fig. 1** with the addition of the central MES-2/MES-6 interacting region; domains/motifs shown are WD-domains (WD40) in MES-6 and LIN-53, and MES-6 binding (MES-6b), Swi3, Ada2, N-CoR and TFIIIB DNA-binding domain 1 like (SANT1), Motif connecting SANT1 and SANT2 (MCSS), SANT2, CXC, and the Su(var)3-9, EZ and Trx domain (SET) in MES-2^26^. We note that the region around the potential MCSS/SANT2 domains in MES-2 is substantially diverged compared with EZH2, yet still displays considerable structural similarity.

Our sequence and structural similarity searches, however, were not able to detect the Zn domain in MES-3 (**Fig. 1b)**, which is normally one of the easiest to identify domains. The absence of Zn is unanticipated as Zn and ZnB in SUZ12 form an intramolecular contact that interact with the accessory PRC2 subunit JARID2^8^. JARID2 contributes PRC2 targeting in embryonic stem cells^8,27^, and even though SUZ12 Zn is not required for methyltransferase activity *in vitro*, Zn is required for PRC2 nucleosome binding *in vivo*, likely by mediating SUZ12-JARID2 interactions^8,21,27^. Thus, while the contact surface of SUZ12 with the core subunits EZH2/1 and EED seems highly conserved in MES-3, MES-3 lacks elements that mediate interactions to accessory subunits that characterize the two PRC2 sub-complexes in mammals.

We conclude that MES-3, even though diverged, structurally resembles SUZ12 in two large regions that are involved in mediating EZH2/1, EED, and RBBP4/7 binding. It is therefore conceivable that, similarly to SUZ12^7,8^, MES-3 is critical in assembling and maintaining a functional PRC2. The here uncovered sequence and structural similarities as well as the peculiar complementary phylogenetic profiles strongly suggest that MES-3 and SUZ12 are in fact orthologs, albeit that MES-3 has undergone rapid sequence divergence and loss of crucial amino acid motifs as well as the Zn domain. Further *C. elegans* specific evolution of the PRC2 assembly and architecture is likely to also play a role. The PRC2 catalytic lobe, which consist of the SUZ12 VEFS domain in association with EZH2 and EED^7,8^, appears the most structurally conserved part of *C. elegans* PRC2. The most notable differences between SUZ12 and MES-3 reside in the N-terminal targeting lobe, which mediates interaction with RBBP4/7, nucleosomes, and accessory proteins^7,8^. From flies to humans, distinct PRC2.1 and PRC2.2 sub-complexes can be distinguished that differ in associated accessory proteins and have specialized functions^3,25,26,28^. For example, the accessory proteins JARID2 and AEBP3 form part of PRC2.2 and mediate interaction with H2AK119ub1^25^, the product of the PRC1 E3 ubiquitin ligase complex^3^. While homologs of JARID2 and other accessory proteins remain to be identified in *C. elegans*, the reported candidate PRC1 components are not required for germline development, in contrast to PRC2^29^. This may explain the lack of conservation of the Zn domain, which in SUZ12 forms part of the JARID2 interaction surface^8^. Additional characterizations of *C. elegans* PRC2 and its accessory proteins will be needed to further substantiate this hypothesis.

The here described similarities and differences between SUZ12 and MES-3 should facilitate further experiments to elucidate the specific mechanisms by which MES-3 acts in PRC2 in *C. elegans*. Our work joins a rapidly growing set of *in silico* predictions of previously undetected homologies made possible by unprecedented advances in deep-learning driven structure prediction^22,30^.

## LIMITATION OF THE SUDY

We capitalized on recent advantages on computational prediction approaches that enable to derive high-quality structures of protein monomers or multimers^23,24^, which enables to study protein function and evolution at unprecedented scale^22,30^. We demonstrate that MES-3 is a diverged ortholog of SUZ12, and that MES-3 may associate with MES-2, MES-6, and LIN-53, similar to the orthologous proteins in human PRC2. However, this study is strictly based on computational predictions, and thus further experiments will be needed to support our predictions and to elucidate how MES-3 functions in *C. elegans* PRC2. This may come, for instance, from resolving the structure of PRC2 in *C. elegans* as well as from genetic engineering experiments of MES-3 in which predicted conserved amino acids and interaction surfaces are modulated, in combination with biochemical and phenotypic characterization.

## ACKNOWLEDGEMENTS

We would like to thank Danny Hancock for the constructing the phylogenetic profiles of PRC2 core members.

## AUTHORS CONTRIBUTION

B.S., S.v.d.H, and M.F.S. conceived the study, performed the experiments, analyzed the data, and drafted the manuscript.

## STARS METHODS

### Resource availability

#### Lead contact

- Further information and requests for resources and data should be directed to and will be fulfilled by the lead contact, Michael F. Seidl.

#### Materials availability

- This study did not generate raw sequencing data.
- Sequence data and software is publicly available as of the date of publication, and accession numbers and software are listed in the key resources table.
- Any additional information required to reanalyze the data reported in this paper is available from the lead contact upon request.

### Method details

#### Sequence similarity searches

We predicted the occurrence of orthologous sequences of the PRC2 core components in diverse Metazoans based on previously computed ortholog assignments from Orthofinder^31^, Broccoli^32^, EggNOG^33^, and SonicParanoid^34^ on a set of reference animal genomes^35^. We manually inspected these orthology assignments based on consistency, which was further corroborated as the predicted occurrences of PRC2 subunits inferred from our assignments consistently matched those published previously (e.g., ref^36,37^).

For sensitive profile-vs-profile searches, we used HHPRED as provided on the MPI Bioinformatics Toolkit server^38^. We performed one search using *C. elegans* MES-3 (uniport: Q10665; MES3_CAEEL) as query and profiles of the human proteome as database, which found as best hit the human SUZ12 protein (ncbi:NP_056170.2) with an e-value 860 and score 38.4. Next, a reciprocal search was performed with human SUZ12 as query and the *C. elegans* proteome as database, which found as best hit MES-3 with an e-value of 970 and score of 28.7; human SUZ12 and *C. elegans* MES-3 are thus in a reciprocal best hit relation of sequence profiles, which is a clear indication for orthology^39^.

#### SUZ12 and MES-3 structure prediction and comparison

We predicted the protein structures of SUZ12 (uniprot:Q15022) and MES-3 (uniprot:Q10665) using a local Alphafold2^23^ instance (version 2.1; five monomer models^23^ as well as model 2 ptm^23^ to obtain the predicted aligned errors, full genetic database, and maximum template date: 01-11-2021). We compared the here predicted with the experimentally determined (rcsbpdb:6WKR-A^25^) structure of SUZ12 using the sequence-independent structure comparisons with super, which is implemented in pymol. Motifs in SUZ12 were selected based on amino acid coordinates^26^ (amino acid coordinates are shown in **Fig. 1b**), and extracted from pdb files using pdb-tools^40^; extracted motifs and domains were subsequently structurally imposed onto the predicted MES-3 using super and/or cealign on the C-alpha atoms, and the root mean square deviation (RMSD; presented in Å) between the structures was used as a measure of structural divergence; an RMSD below 2 Å is generally considered to indicate two very similar structures. We furthermore used TM-align (version 20190822; default parameters)^41^ as well as Dali^42^ to obtain sequence-independent structure alignments between SUZ12 and MES-3 (sub)structures; TM align TM-scores 0.5 < x < 1 and Dali Z-scores > 2 typically indicate similar folds. Disordered regions in the protein sequences were predicted using IUPRed3 (default settings)^43^. The protein (sub)structures were visualized using pymol, and the data visualization was performed with python seaborn.

#### PRC2 complex structure prediction and comparison

We predicted the monomeric structures of the members of the PRC2 core complex, MES-2 (EZH2; uniprot:O17514), MES-6 (EED; uniprot:Q9GYS1), and LIN-53 (RBBP4; uniprot:P90916), with Alphafold2 and compared these monomeric predictions with experimentally predicted structure of human PRC2 members (rcsbpdb:6WKR^25^) as described above. We predicted multi-chain PRC2 complex interactions of MES-2, MES-3, and MES-6 as well as MES-3 and LIN-53 using Alphafold2-multimer^24^ (version 2.2; five multimer models^24^ with each five seeds, full genetic database, and maximum template date: 01-11-2021). Predicted multimer models were compared with monomer models using super as well as TM-align^41^ as described above, and interaction interfaces between protein pairs within complexes were predicted using pymol (default settings).

