## Supplementary Figures S1-2 for "*Caenorhabditis elegans* MES-3 is a highly divergent ortholog of the canonical PRC2 component SUZ12"

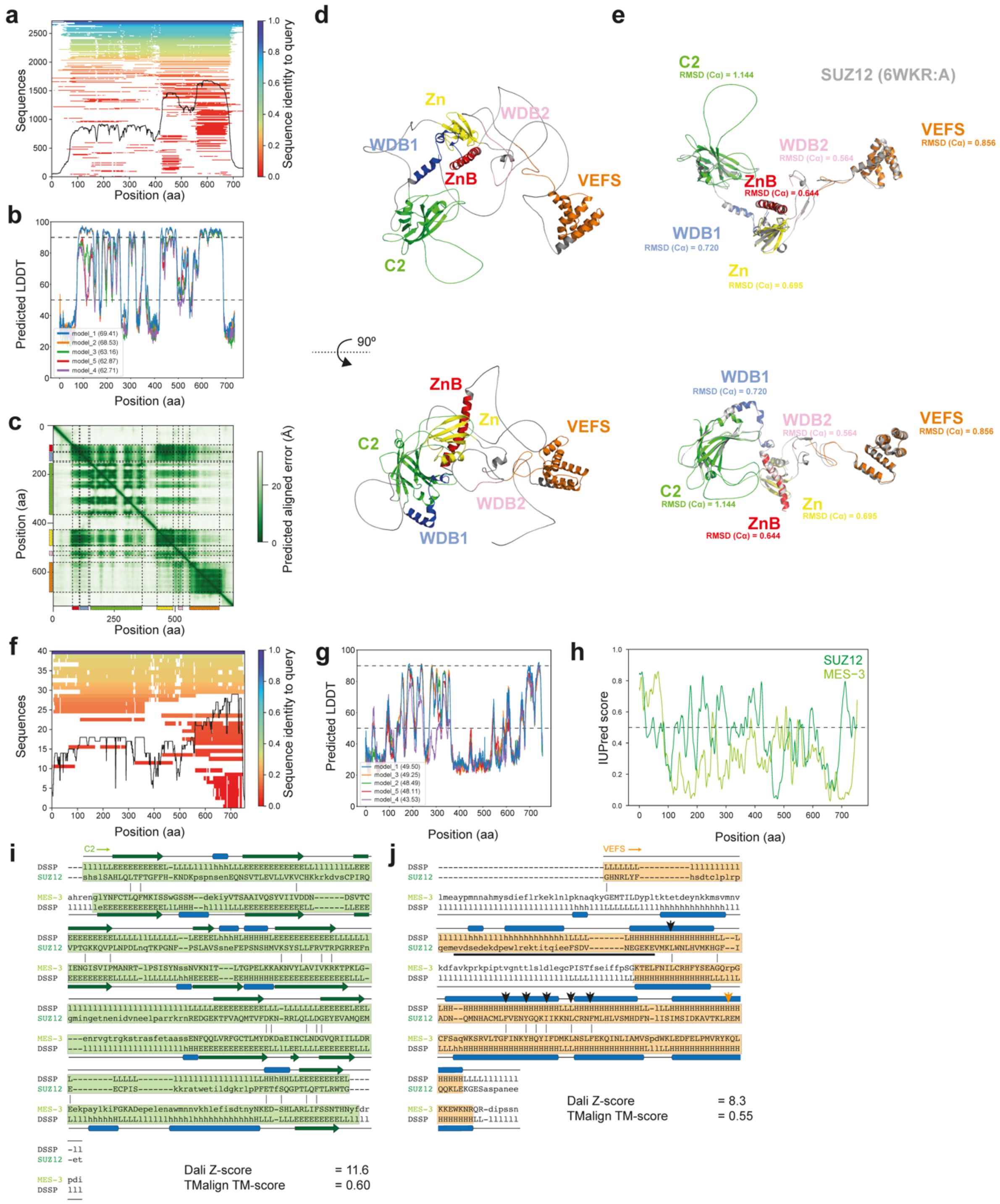


**Figure S1. SUZ12 and MES-3 share significant structural similarity, related to Figure 1**

**a.** The number of homologous protein sequences per position of SUZ12 retrieved from different genetic databases used by Alphafold2[1]. We recovered in total 2,720 (deduplicated) sequences that contributed to the multiple sequence alignment as well as 14 template structures. **b.** The predicted Local Distance Difference Test (pLDDT) per position for each of the five Alphafold2 predicted models of SUZ12 as well as the average pLDDT per model is shown, indicating that the individual six domains and motif of SUZ12 are predicted with high confidence; regions with pLDDT < 50 can be interpreted as disordered regions[1], and quality thresholds of 50 (low confidence) and 90 (high confidence) are indicates as dashed lines. **c**. The predicted aligned error (in Å; expected position error at residue x when aligned to position residue y; based on model 2 ptm) of the predicted SUZ12 structure indicates that the Alphafold2 prediction is confident about the relative position of most domains and motifs except for the C-terminal of the VEFS domain. The six domains and motifs are shown along the x- and y-axis; the zinc finger binding (ZnB; red), WD-domain binding 1 (WDB1; blue), C2 domain (green), zinc finger (Zn; yellow), WD-domain binding 2 (WDB2; pink), and VEFS (orange). **d**. The predicted Alphafold2[1] protein structure of SUZ12 displays multiple globular regions, and the six domains and motifs are shown by different colors, similar to **c**. **e.** The six domains and motifs extracted from the predicted SUZ12 structure are superimposed on the experimentally determined structure of SUZ12 (rcsbpdb:6WKR-A[2]; grey; note, the experimentally determined structure does not cover the entire protein sequence and contains unresolved regions). The root-mean-square deviation (RMSD) between the predicted and experimentally determined SUZ12 (sub)structures is indicated; RMSD < 2 Å indicate structures with very similar fold. **f**. The number of homologous protein sequences per position of MES-3 retrieved from different genetic databases used by Alphafold2[1]. As expected (**Fig. 1a**), compared with SUZ12 we only identified 40 homologous sequences that were included into the structure prediction. **g**. The predicted Local Distance Difference Test (pLDDT) per position for each of the five Alphafold2 predicted models of MES-3 as well as the average pLDDT per model is shown, indicating that part of MES-3 is globular and predicted with high confidence (pLDDT > 80). **h.** Prediction of disordered regions for SUZ12 (dark green) and MES-3 (light green). MES-3 is predicted to contain a largely disordered N-terminal region (< ~80 amino acids), while most of the protein is predicted to have reduced disorder, which coincides with high-confidence prediction in panel **g**. **i**. Sequence-independent structure alignment of the C2 regions of SUZ12 (aa 150 - 365) and MES-3 (aa 145 - 370) reveals significantly structural similarity (Dali Z-score = 11.6; TM-score = 0.6), especially along the beta sheets. The secondary structural inference by DSSP[3] is displayed (L/l, loop; H/h, helix; E/e, sheet; matching structures are indicated in capital). **j**. Sequence-independent structure alignment of the VEFS regions of SUZ12 (aa 560 - 700) and MES-3 (aa 560 - 754) reveals significantly structural similarity

(Dali Z-score = 8.3; TM-score = 0.55), especially along the alpha helices in the C-terminus; a region previously shown to stimulate histone methyltransferase activity in SUZ12[4] (pos. 580 to 612) is highlighted by a black bar, and individual amino acids important for PRC2 assembly[4] are shown by black arrows. The annotated VEFS domain (**Fig. 1b;** aa 560 – 682; orange arrow) is shorter than inferred by the Alphafold2 (aa 560 - 690), based on the occurrence of a conserved alpha helix between SUZ12 and MES-3. The secondary structural inference by DSSP[3] is displayed along the sequences (L/l, loop; H/h, helix; E/e, sheet; matching structures are indicated in capital).

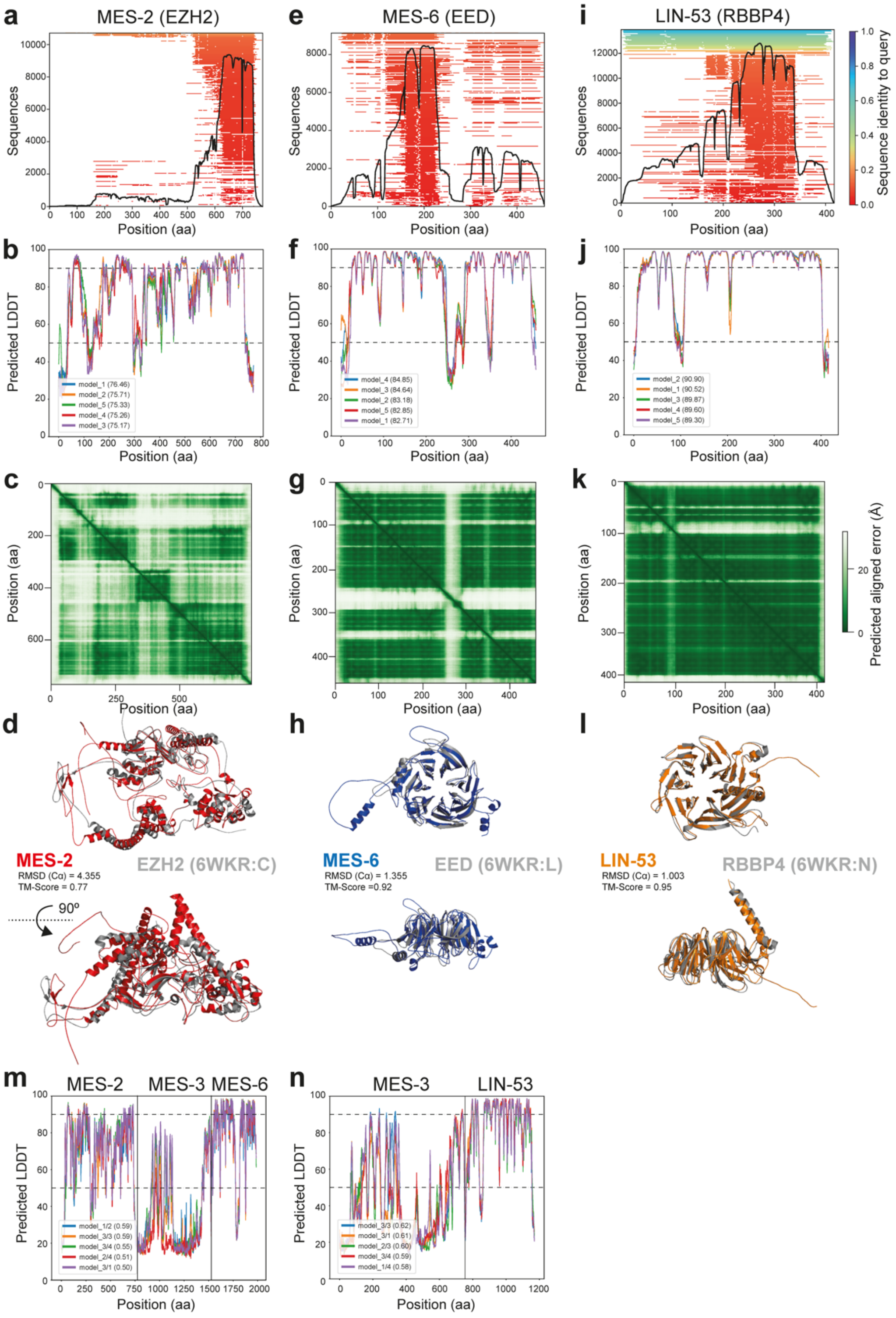


**Figure S2. Structural prediction of *C. elegans* PRC2 complex components, related to Figure 2**

The number of homologous protein sequences per position of MES-2 (EZH2; **a**), MES-6 (EED; **e**), and LIN-53 (RBBP4; **i**) retrieved from different genetic databases used by Alphafold2[1]. We recovered in total 10,712, 9,126, and 13,899 (deduplicated) sequences, respectively, that contributed to the multiple sequence alignment as well as the maximum number (20) of template structures. The predicted Local

Distance Difference Test (pLDDT) per position for each of the five Alphafold2 models of MES-2 (**b**), MES-6 (**f**), and LIN-53 (**j**) as well as the average pLDDT per model is shown, indicating that the proteins are predicted with high confidence; regions with pLDDT < 50 can be interpreted as disordered regions[1], and quality thresholds of 50 (low confidence) and 90 (high confidence) are indicates as dashed lines. The predicted aligned error (in Å; expected position error at residue x when aligned to position residue y; based on model 2_ptm) of the predicted MES-2 (**c**), MES-6 (**g**), and LIN-53 (**k**) structure indicates that the Alphafold2 prediction is confident about the relative position of the domains and motifs. The predicted MES-2, MES-6, and LIN-53 structures were superimposed on the experimentally determined structure of EZH2, EED, and RBBP4, respectively (rcsbpdb:6WKR[2]); the experimental structures are shown in grey and the predicted *C. elegans* structures according to **Fig. 1b**. The root-mean-square deviation (RMSD) and TM-Score between the predicted and experimentally determined structures is indicated; RMSD < 2 Å and TM align TM-scores 0.5 < x < 1 indicate structures with very similar fold. The predicted Local Distance Difference Test (pLDDT) per position for the top five Alphafold2-multimer models (model and seed) of the MES-2, MES-3, and MES-6 (**m**) and the MES-3 and LIN-53 complexes (**n**) as well as the weighted predicted (interface) TM-score per model is shown.
